# Loss of histone H3.3 results in DNA replication defects and altered origin dynamics in *C. elegans*

**DOI:** 10.1101/854455

**Authors:** Maude Strobino, Joanna M. Wenda, Florian A. Steiner

## Abstract

Histone H3.3 is a replication-independent variant of histone H3 with important roles in development, differentiation and fertility. Here we show that loss of H3.3 results in replication defects in *Caenorhabditis elegans* embryos. To characterize these defects, we adapt methods to determine replication timing, map replication origins, and examine replication fork progression. Our analysis of the spatiotemporal regulation of DNA replication shows that despite the very rapid embryonic cell cycle, the genome is replicated from early and late firing origins and is partitioned into domains of early and late replication. We find that under temperature stress conditions, additional replication origins become activated. Moreover, loss of H3.3 results in impaired replication fork progression around origins, which is particularly evident at stress-activated origins. These replication defects are accompanied by replication checkpoint activation, a prolonged cell cycle, and increased lethality in checkpoint-compromised embryos. Our comprehensive analysis of DNA replication in *C. elegans* reveals the genomic location of replication origins and the dynamics of their firing, and uncovers a role of H3.3 in the regulation of replication origins under stress conditions.

## Introduction

At every cell cycle, the entire genome is replicated exactly once for accurate transmission of genetic information to the daughter cells. To ensure rapid and uniform duplication of the genome, DNA replication is initiated in a bi-directional way at many different sites in the genome, called replication origins. To prevent re-replication and to control the activity of origins under replication stress, replication initiation is composed of two non-overlapping steps called ‘origin licensing’ and ‘origin firing’. Origin licensing occurs during G1 through the formation of the pre-replicative complexes. During this process, the origin recognition complex (ORC) binds to the origins and recruits the replication licensing factors Cdt1 and Cdc6, which together load the complex of the minichromosome maintenance proteins 2-7 (Mcm2-7), a helicase that unwinds DNA ahead of the replication fork (Sonneville et al., 2012). To ensure that each origin fires only once per cell cycle, Cdt1 and Cdc6 are subsequently removed from the nucleus or degraded to prevent the Mcm2-7 complex from rebinding origins that have already fired (Blow and Dutta, 2005; Feng and Kipreos, 2003; Kim and Kipreos, 2007). Origin firing occurs upon entry into S phase, through binding of different initiation factors like Sld2 (RecQL4/RecQ4 in humans), Sld3 (Treslin/TICRR in humans) and Dpb11 (TopBP1 in humans), and phosphorylation of the Mcm2-7 complex (Bruck and Kaplan, 2015; Kanemaki and Labib, 2006; Labib and Diffley, 2001).

These firing factors are present in limiting amounts (Mantiero et al., 2011; Tanaka et al., 2011), and all origins therefore do not fire synchronously at the start of S phase. Instead, some origins fire early and some late, resulting in large domains of early and late replication (Desprat et al., 2009; Dileep and Gilbert, 2018; Farkash-Amar et al., 2008; MacAlpine et al., 2004; Takahashi et al., 2019). These domains may change during development, and it has been shown in multiple organisms that early replicating domains are associated with transcriptionally active regions (Hiratani et al., 2010, 2008; Schübeler et al., 2002; Therizols et al., 2014; White et al., 2004; Woodfine et al., 2004).

Upon replication origin firing, the replication forks move bi-directionally until replication bubbles of opposing directions meet and the replication of the corresponding stretch of chromosome is complete. Under specific conditions, e.g. conflicts between transcription and replication forks, depletion of nucleotides, or depletion of histones, replication forks can stall or collapse (Bester et al., 2011; Groth et al., 2007; Macheret and Halazonetis, 2018). This replication stress can lead to DNA damage (Macheret and Halazonetis, 2015; Técher et al., 2017), and triggers the activation of the ATR checkpoint, which phosphorylates the Chk1 kinase, leading to cell cycle arrest (Hyun et al., 2016; Liu et al., 2000; Moiseeva et al., 2019). As a response to moderate replication stress, cells can activate additional origins (Técher et al., 2017). These origins, called latent or dormant origins, are inactive and do not fire during the undisturbed S phase, but act as backup origins in case of replication stress (Boos and Ferreira, 2019; Woodward et al., 2006). How firing origins are selected over dormant origins, and under what conditions dormant origins are activated is unclear (Fragkos et al., 2015).

It is widely accepted that replication origins are not defined by DNA sequences in higher eukaryotes and therefore cannot be predicted from genome sequences, making their identification challenging. Numerous assays have been developed to map replication origins in different organisms (Prioleau and MacAlpine, 2016), mostly by mapping different replication elements such as nascent DNA strands (Besnard et al., 2012), replication bubbles (Mesner et al., 2013), Okazaki fragments (Petryk et al., 2016), replication initiation sites (Langley et al., 2016), or initiation factors (Dellino et al., 2013; Miotto et al., 2016). Despite these efforts, the number, location and dynamic regulation of replication origins remains controversial. For example, two studies in *C. elegans* recently provided genome-wide maps of replication origins using two different methods, and arrived at largely different conclusions (Pourkarimi et al., 2016; Rodríguez-Martínez et al., 2017). The first study used Okazaki fragment sequencing to identify ∼2’000 origins and concluded that replication origins are defined prior to the broad onset of zygotic transcription and maintained during development (Pourkarimi et al., 2016). The second study applied mapping of small RNA-primed nascent DNA strands to identify ∼16’000 origins and described a major re-organization of replication origins after gastrulation (Rodríguez-Martínez et al., 2017). Additional approaches will therefore be required to understand how origins are defined and dynamically regulated.

The variant histone H3.3 has been linked to different steps of DNA replication. It is enriched at replication origins in *Drosophila melanogaster* and *Arabidopsis thaliana* (Deal et al., 2010; Eaton et al., 2011; MacAlpine et al., 2010; Paranjape and Calvi, 2016; Stroud et al., 2012), and loss of H3.3 results in a slower replication fork in chicken cells (Frey et al., 2014). The association of H3.3 with early replication domains, and its recruitment to sites of DNA repair have also been described in human cells (Adam et al., 2013; Clément et al., 2018). H3.3 is incorporated into transcriptionally active regions as well as in centromeric and telomeric repeats (Buschbeck and Hake, 2017; Henikoff and Smith, 2015), but the link between its genomic location and its function is not always clear. Loss of H3.3 results in lethality or sterility in most organisms (Hödl and Basler, 2009; Jang et al., 2015; Sakai et al., 2009; Tang et al., 2015; Wollmann et al., 2017). The *C. elegans* genome contains five genes encoding H3.3 homologues, and we recently showed that mutation of all five genes resulted in viable worms with a reduced number of viable offspring at higher temperatures (Delaney et al., 2018).

Given the role of H3.3 in DNA replication in other organisms, we speculated that the reduced brood size observed in *C. elegans* H3.3 null mutants is caused by defects in DNA replication. Here we show that at higher temperatures, loss of H3.3 causes mild DNA replication defects and activation of the embryonic DNA replication checkpoint. To investigate these defects, we adapt ChEC-seq, Repli-seq, EdU-seq and DNA combing for use in *C. elegans*, which we apply to define and characterize replication timing and replication origins. We find that during S phase, replication and the firing of replication origins are temporally regulated. H3.3 is enriched at replication origins, but is not required for origin firing. Instead, loss of H3.3 results in delayed fork progression around replication origins, in particular around origins that are only active under temperature stress conditions. Together, our results uncover a role of H3.3 in DNA replication in *C. elegans* and show that replication origin use can be adapted in response to stress.

## Results

### Genome-wide replication timing in *C. elegans* embryos

Histone H3.3 has been linked to the regulation of DNA replication in a variety of organisms (Adam et al., 2013; Clément et al., 2018; Deal et al., 2010; Eaton et al., 2011; Frey et al., 2014; MacAlpine et al., 2010; Paranjape and Calvi, 2016; Stroud et al., 2012). We therefore hypothesized that the reduced brood sizes observed in *C. elegans* H3.3 null mutants under mild temperature stress are caused by defects in DNA replication (Delaney et al., 2018). Our first aim was to determine whether loss of H3.3 causes global changes in the temporal program of genome replication. To characterize replication timing during the rapid cell cycle in early *C. elegans* embryogenesis, we adapted Repli-seq for use in this organism (Hansen et al., 2010). This method relies on the incorporation of the thymidine analogue 5-ethynyl-2’-deoxyuridine (EdU) into newly replicated regions of the genome (Figure 1A). To separate early and late replicated genomic regions, we pulse-labelled asynchronous embryonic cell populations with EdU, then sorted the cells according to their DNA content and sequenced EdU-containing fragments. We obtained cells that just entered S phase (early replication), showed ongoing DNA synthesis (mid replication), or were at or near completion of replicating the genome (late replication) (Figure S1A). These Repli-seq experiments revealed that the genome is partitioned into mutually exclusive domains of early and late replication (Figure 1B, C). Surprisingly, different chromosomes display distinct overall replication timing patterns. Domains of early replication are enriched on chromosomes I, II and III, while domains of late replication are more prevalent on chromosomes IV, V and X (Figure 1C and Figure S1B). We found that domains of early replication correlate well with the presence of HIS-72, the most highly expressed *C. elegans* H3.3 homologue (Figure 1B and Figure S1C). However, H3.3 null mutant worms showed largely the same early and late replication domains as wild-type worms, indicating that H3.3 is not a major driver of global DNA replication timing (Figure 1B, C and Figure S1B). Genes in domains of early replication show higher levels of expression in embryo compared to genes in domains of later replication, consistent with the observation from other organisms that regions of open chromatin replicate early (Ekundayo and Bleichert, 2019; Prioleau and MacAlpine, 2016) (Figure 1D). However, there is no enrichment of large genes in late replicating domains that would suggest potential interference between transcription and replication under stress conditions, as observed in human cells (Figure 1E) (Macheret and Halazonetis, 2018).

**Figure 1.**
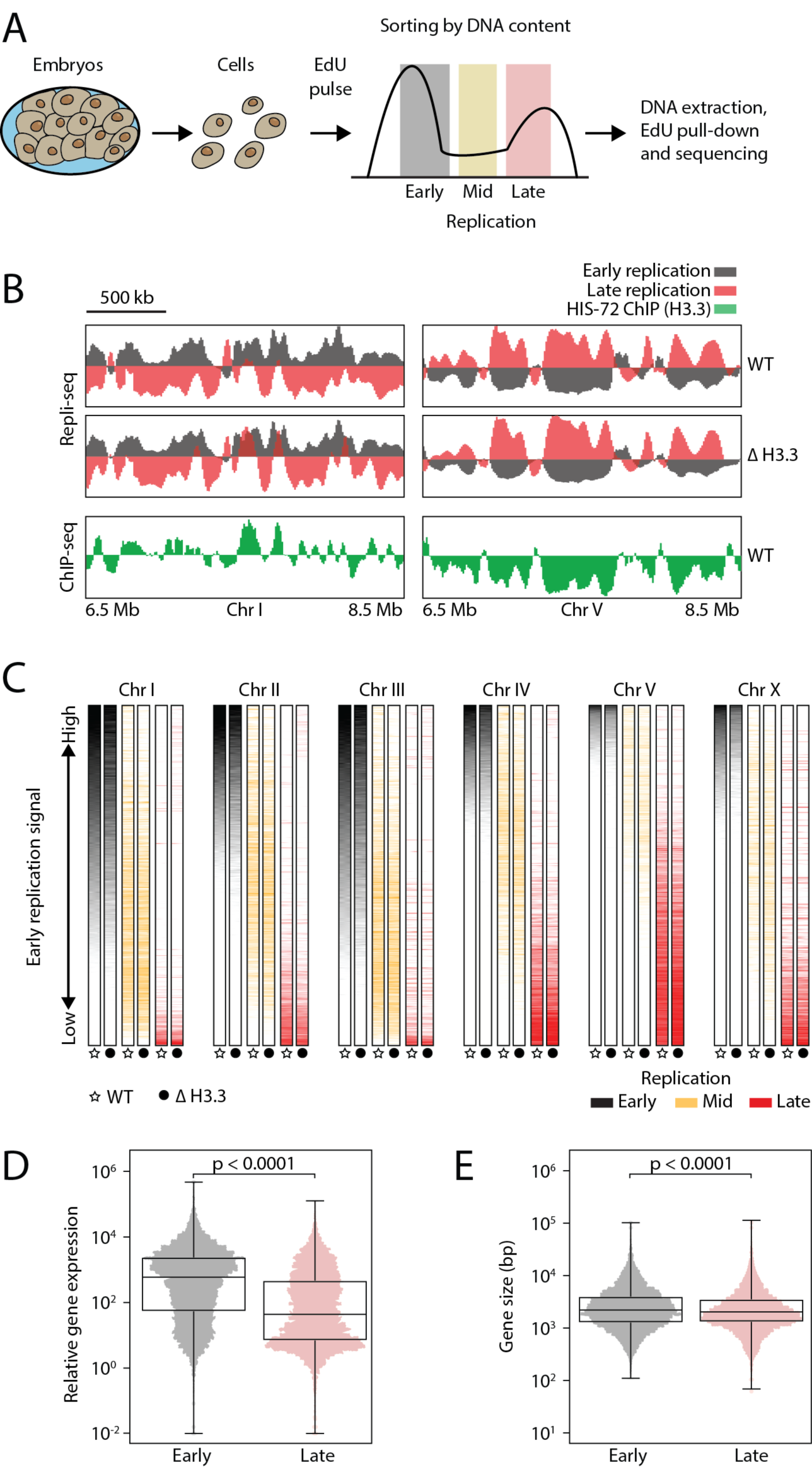
Replication timing is unchanged in absence of H3.3. (**A**) Schematic description of Repli-seq. Embryonic cells were dissociated, exposed to an EdU pulse for 10 min and sorted according to their DNA content. EdU-labelled DNA was sequenced and mapped to the genome. (**B**) Representative genome browser views of Repli-seq and H3.3 ChIP-seq. Repli-seq signal is shown for wild type (WT) and H3.3 null mutant (Δ H3.3) worms on regions of chromosomes I and V. Early S phase is shown in black and late S phase in red. HIS-72 (H3.3) ChIP-seq signal is shown in green for the same regions. ChIP-seq data from (Delaney et al., 2019). (**C**) Color-coded replication timing for each chromosome. Repli-seq signal from early (black), mid (orange) and late (red) S phase for WT and Δ H3.3. Data for each chromosome was sorted according to the signal of early S phase in WT. (**D**) Gene expression levels for the genes present in domains of early and late replication. RNA-seq data from (Kramer et al., 2015). (**E**) Genes sizes for the genes present in domains of early and late replication.

### Mapping replication origins by ChEC-seq and EdU-seq

Given the presence of domains of early and late replication, we reasoned that there should be distinct populations of early and late firing replication origins. With the enrichment of H3.3 at replication origins in other organisms, we speculated that loss of H3.3 may cause subtle defects in the position or timing of origin firing in H3.3 null mutants. We therefore next aimed to define replication origins and their timing during S phase. Previous origin mapping approaches in *C. elegans* do not allow for the analysis of origin dynamics during the cell cycle (Pourkarimi et al., 2016; Rodríguez-Martínez et al., 2017). We therefore took an alternative approach that combines the mapping of replication origin factors and the observation of nucleotide incorporation at origins. We first profiled the genomic binding sites of proteins involved in replication origin licensing and firing. We selected two conserved proteins, CDT-1 and TRES-1, that are involved in different steps of origin activation (Figure 2A). CDT-1 is the homologue for human Cdt1 and is implicated in the licensing of all origins during G1 phase (Zhong et al., 2003) (Figure 2A). TRES-1 is the homologue of human Treslin and is subsequently recruited exclusively to the origins that are going to fire (Guo et al., 2015; Kumagai et al., 2010). Immunofluorescence microscopy confirmed that CDT-1 is present on chromatin during anaphase and telophase, but levels become undetectable during S-phase. TRES-1 only appears on chromatin during telophase, but remains present during S phase (Figure 2B). These observations are consistent with their roles in origin licensing and origin firing, respectively. Typically, genomic protein binding sites are profiled by chromatin immunoprecipitation. However, this approach depends on antibody availability and solubility of the chromatin component of interest, all of which can be limiting factors. To circumvent these problems, we adapted Chromatin Endogenous Cleavage (ChEC) to *C. elegans*. This method relies on the fusion of the protein of interest with micrococcal nuclease (MNase), which cuts DNA around the binding sites of the fusion protein and thereby releases DNA fragments that can be sequenced and mapped (Schmid et al., 2004; Zentner et al., 2015) (Figure 2C). We generated MNase fusion constructs at the endogenous *cdt-1* and *tres-1* loci using CRISPR/Cas9. MNase insertions resulted in worms that are phenotypically wild-type. To generate genome-wide binding maps of CDT-1 and TRES-1, we performed ChEC-seq experiment on purified embryonic nuclei and identified peaks of CDT-1 and TRES-1 signal. We observed CDT-1 and TRES-1 peaks that are distributed across the genome in both datasets. Moreover, as expected, the CDT-1 peaks mostly overlap with TRES-1 peaks (licensed and firing), but some CDT-1 peaks show no corresponding TRES-1 signal (licensed only) (Figure 2D).

**Figure 2.**
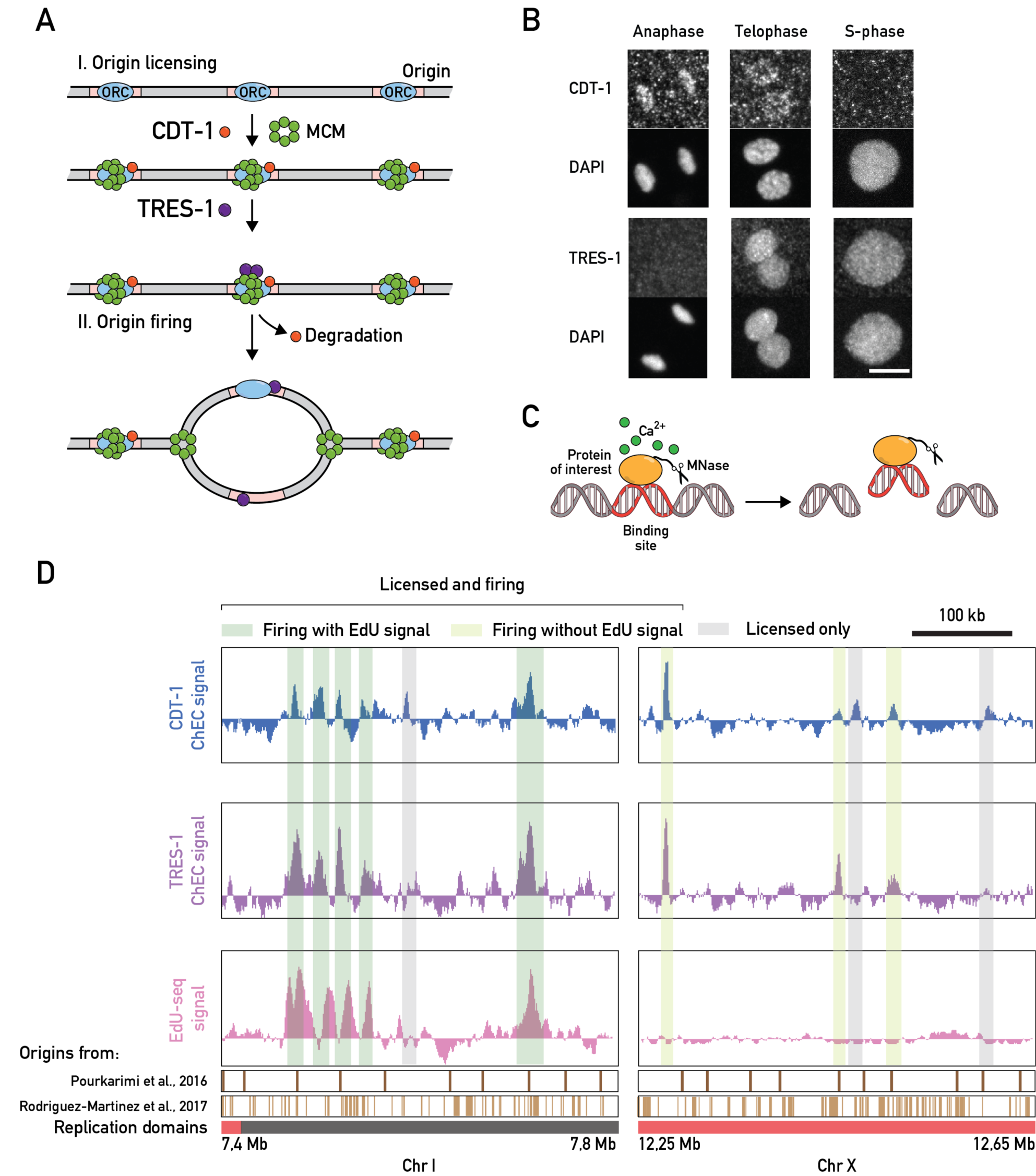
Identification of replication origins in *C. elegans* embryos. (**A**) Schematic description of the roles of CDT-1 and TRES-1 in replication origin firing. CDT-1 is required for the licensing of all origins. TRES-1 is recruited only to origins that fire. (**B**) Localization of HA::CDT-1 and FLAG::TRES-1 during the cell cycle. Immunofluorescence images using anti-HA and anti-FLAG antibodies and DAPI are shown. Scale bar represents 5 µm. (**C**) Schematic description of ChEC-seq. The protein of interest is fused with MNase. Upon activation with calcium in purified nuclei, MNase cleaves and releases DNA fragments at the binding sites of the fusion proteins. These small fragments are isolated and sequenced. (**D**) Representative genome browser views of CDT-1 ChEC-seq (blue), TRES-1 ChEC-seq (violet) and EdU-seq (pink) signal. Identified origins are highlighted in gray (licensed only) or dark and light green (licensed and firing with and without EdU signal, respectively). Domains of late (red) and early (gray) replication and positions of replication origins identified in previous studies are shown below the genome browser.

The CDT-1 and TRES-1 ChEC-seq data allowed us to discriminate between licensed and firing origins, but we could not establish the timing of origin firing within S phase. To identify replication origins that fire immediately after entry into S phase, we modified the Repli-seq protocol. Instead of analyzing asynchronous cell populations, we used hydroxyurea (HU) to synchronize cells at the entry of the S phase. Upon HU treatment, virtually all cells become arrested at the entry of the S phase, and a large part of the cell population proceeds with DNA replication upon removal of HU (Figure S2A). We combined the release into S phase with a second HU block in the presence of EdU, allowing the cells to replicate short stretches of DNA around early replication origins before arresting. The EdU-containing DNA fragments were then isolated and sequenced. These EdU-seq experiments revealed discrete peaks that are located in close proximity to with a subset of the CDT-1 and TRES-1 peaks (licensed and firing with EdU signal) (Figure 2D). The EdU-seq signal sometimes appear as double peaks flanking the CDT-1 and TRES-1 peaks, indicating that we detect short fork movements away from the replication origins. We conclude that the sites enriched for EdU-seq, CDT-1 and TRES-1 signal correspond to replication origins that fire upon entry into S phase.

### Classifications of origins genome wide

To obtain a map of embryonic replication origins genome-wide, we combined the peak calls from the CDT-1 ChEC-seq, TRES-1 ChEC-seq and EdU-seq experiments. In total, we obtained a map of 1110 origins. Based on their signal in the three datasets, unsupervised clustering distinguished three types of origins, which we classified as “early firing”, “late firing” and “dormant” (Figure 3A). Early firing origins show enrichment in all three datasets, as they are licensed (CDT-1) and fire at the onset of S phase (TRES-1 and EdU-seq). Late firing origins show enrichment in CDT-1 and TRES-1 ChEC-seq datasets, but not EdU-seq, indicating that they fire after the onset of S phase. Dormant origins show enrichment only in the CDT-1 ChEC-seq dataset, indicating that they are licensed, but do not fire under standard conditions (Figure 3A). While the enrichment of CDT-1 and TRES-1 overlaps within a narrow window at the replication origins, the distribution of EdU-seq signal is more broad, which likely reflects short stretches of fork progression away from the replication origins after the release from the initial HU block. As expected, firing origins are found mostly outside of gene-coding regions, while dormant origins are enriched within genes (Figure S2B). The genomic localization of the replication origins identified here largely overlap with those previously identified (Figure S2C), but our mapping approach offers the advantage of capturing origin firing dynamics during the cell cycle, and also includes dormant origins that may act as backup origins for use under specific conditions. Similar to *D. melanogaster* and *A. thaliana* (Deal et al., 2010; Eaton et al., 2011; MacAlpine et al., 2010; Paranjape and Calvi, 2016; Stroud et al., 2012), H3.3 is enriched around the origins of all three categories (Figure 3B), suggesting that it may play a role in origin identity or activation.

**Figure 3.**
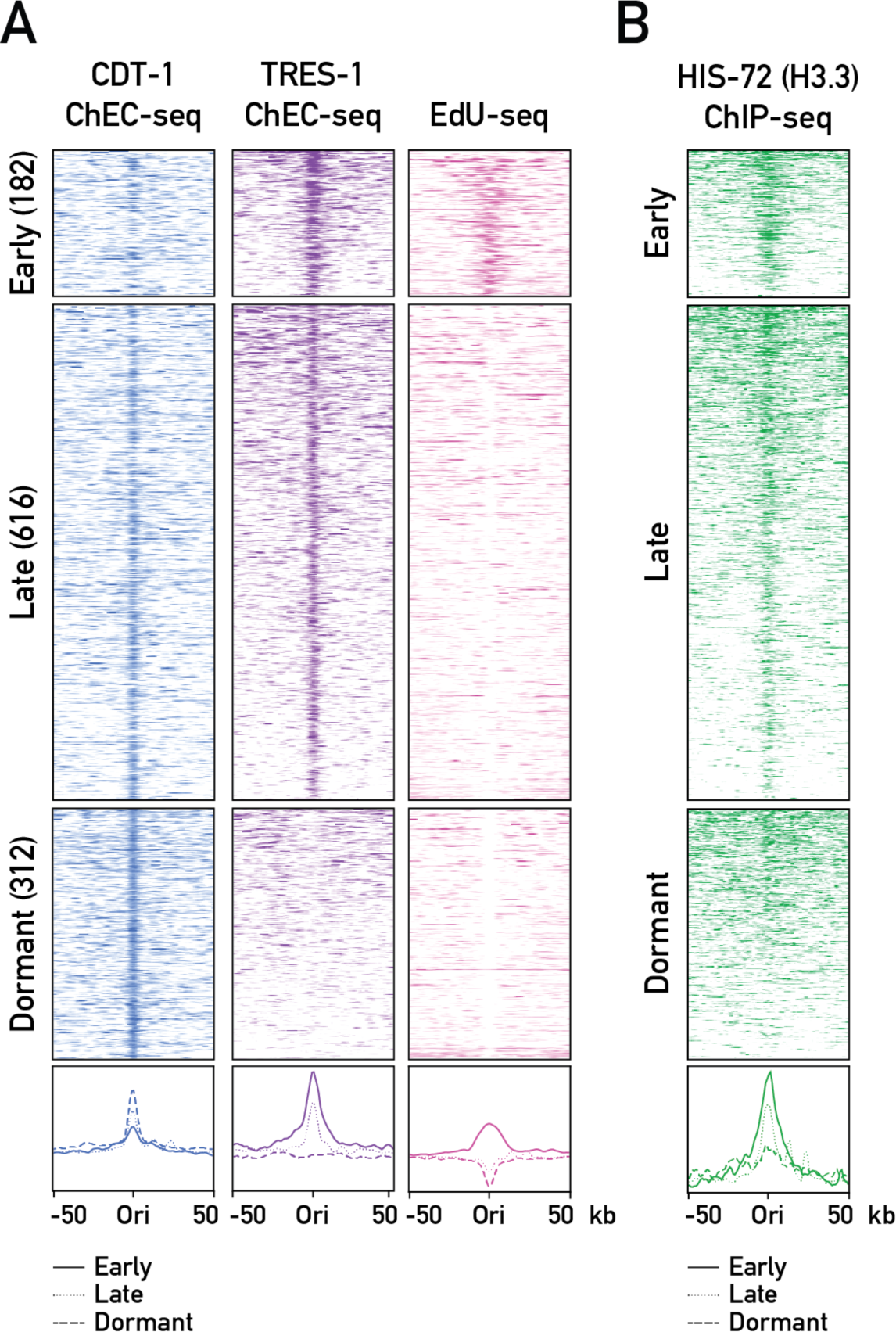
Classification of replication origins in *C. elegans* embryos. (**A**) Heatmaps (top) and average plots (bottom) of CDT-1 ChEC-seq (blue), TRES-1 ChEC-seq (violet) and EdU-seq (pink) signal at replication origins. Origins were identified by peak calling using the CDT-1 ChEC-seq, TRES-1 ChEC-seq and EdU-seq datasets obtained from worms grown at 20°C and separated into early, late and dormant origins through unsupervised clustering. Signal is shown for a 50 kb window around each origin. (**B**) Heatmaps (top) and average plots (bottom) of HIS-72 (H3.3) ChIP-seq signal (Delaney et al., 2019) at replication origins, as in (A).

### Altered origin dynamics in H3.3 null mutants

We next compared the distribution of firing origins in wild type and H3.3 null mutant worms. For this, we repeated the TRES-1 ChEC-seq and the EdU-seq experiments in wild type and H3.3 null mutant strains at 25°C, where a difference in brood-size between the two strains is observed. The TRES-1 ChEC-seq experiments showed that most origins licensed in wild type worms were also utilized in absence of H3.3 (Figure 4A). In both strains, we also observed weak TRES-1 signal at origins that are dormant at 20°C. These origins were initially identified solely based on the presence of CDT-1 under normal conditions, and the presence of TRES-1 at 25°C shows that these are functional origins that become activated at higher temperatures. This result confirms the presence of “backup” origins that are inactive during undisturbed S phase, but are activated under replication stress (Boos and Ferreira, 2019; Woodward et al., 2006). We did not detect any difference between the firing of early origins in wild type and H3.3 null mutant based on EdU-seq signal (Figure 4C, 0min). These results indicate that the origin distribution and the firing of early origins remains unchanged upon loss of H3.3.

**Figure 4.**
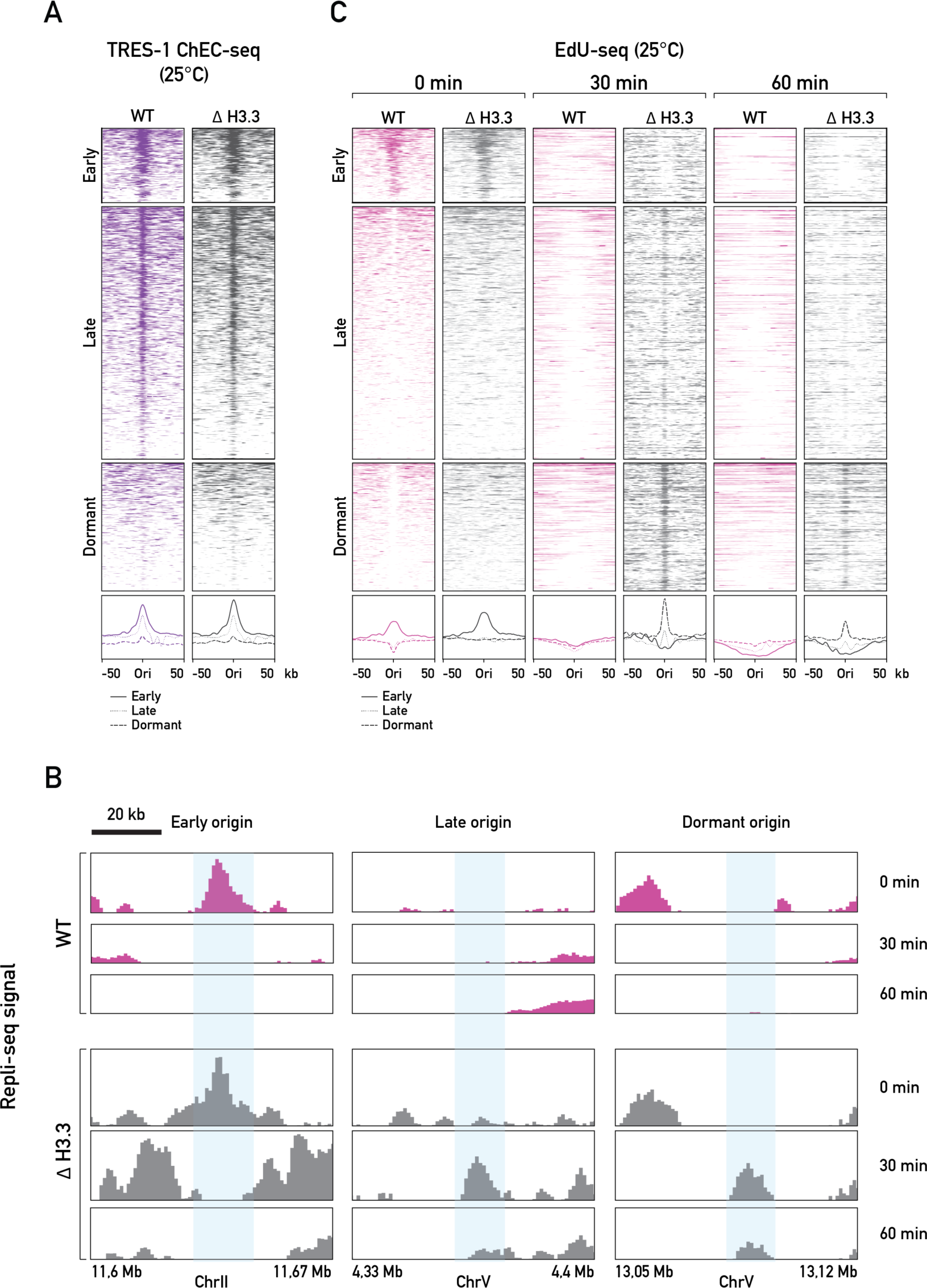
Replication origin dynamics is altered H3.3 null mutants. (**A**) Heatmaps (top) and average plots (bottom) of TRES-1 ChEC-seq signal at replication origins, for wildtype (WT, violet) and H3.3 null mutant (Δ H3.3, gray) worms grown at 25°C. Signal is shown for a 50 kb window around each origin. (**B, C**) EdU-seq time course, for WT (pink) and Δ H3.3 (gray) worms grown at 25°C. (**B**) Genome browser views of EdU-seq signal at 0, 30 and 60 minutes, showing fork progression at representative examples of early, late and dormant origins. (**C**) Heatmaps (top) and average plots (bottom) of EdU-seq signal at 0, 30 and 60 minutes, at all origins. Signal is shown for a 50 kb window around each origin.

Although the firing of early origins is not affected by the loss of H3.3, we were wondering if fork progression following origin firing could be altered. The EdU-seq method not only allows one to identify early firing origins, but can also be used to visualize fork progression around these origins, by adding an EdU pulse at 0, 30 or 60 minutes after the release from the HU block (Figure 4B-C). As expected, the EdU signal enrichment at the early firing origins is no longer present after 30 minutes, and regions occupied by early firing origins incorporate very little EdU after 60 minutes, which reflects the bi-directional movement of the replication forks away from the origins (Figure 4B-C, WT). This bi-directional fork movement is difficult to observe for late origins, as we have no means to synchronize their firing. Nevertheless, the EdU-seq signal at late origins appears depleted at 30 minutes, suggesting that they fire within the first 30 minutes after S-phase onset. In H3.3 null mutants, as mentioned above, early origins appear to fire normally, and we observe bi-directional fork movement. However, we found that the EdU signal remains higher around early origins after 30 and 60 minutes, indicating that fork progression is delayed (Figure 4B-C, Δ H3.3). This delay in fork progression appears at origins of all classes and results in an enrichment of EdU incorporation at late origins. The difference between wild type and H3.3 null mutant cells is even more pronounced at origins that are dormant under standard conditions. These origins fire in both strains at 25°C, as evidenced by the increased presence of TRES-1 compared to 20°C (Figure 3A and Figure 4A), but fork progression from the origins is impaired in H3.3 null mutants, resulting in an accumulation of EdU-seq signal at later time points. The enrichment at late and dormant origins is observed within a very narrow window, suggesting that the forks stall or collapse immediately after origins fire. The coincidence of this EdU-seq signal enrichment with the center of late and dormant origins is particularly remarkable, as these origins were identified solely based on a completely independent method, namely the mapping of the CDT-1 and TRES-1 binding sites. The aberrant fork progression in H3.3 null mutant worms is only seen under mild temperature stress conditions, and is not observed at 20°C (Figure S3), indicating that H3.3 is required for regulating origin dynamics under stress.

### Fork speed is unaffected by loss of H3.3

To establish if loss of H3.3 resulted in a general slowing of the replication fork speed along the chromatin fiber, we adapted DNA combing to *C. elegans (Michalet et al., 1997)*. This method relies on the pulse labeling of DNA by the subsequent incorporation of two thymidine analogues, IdU and CldU. By visualizing the length of IdU and CldU incorporation on stretched individual chromatin fibers, replication fork speed can be estimated (Figure 5A). We found that fork speed is on average 1.4 kb/min (Figure 5B), but did not detect any significant difference between wild type and H3.3 null mutant embryos, indicating that H3.3 is not required for normal progression of the moving replication fork.

**Figure 5.**
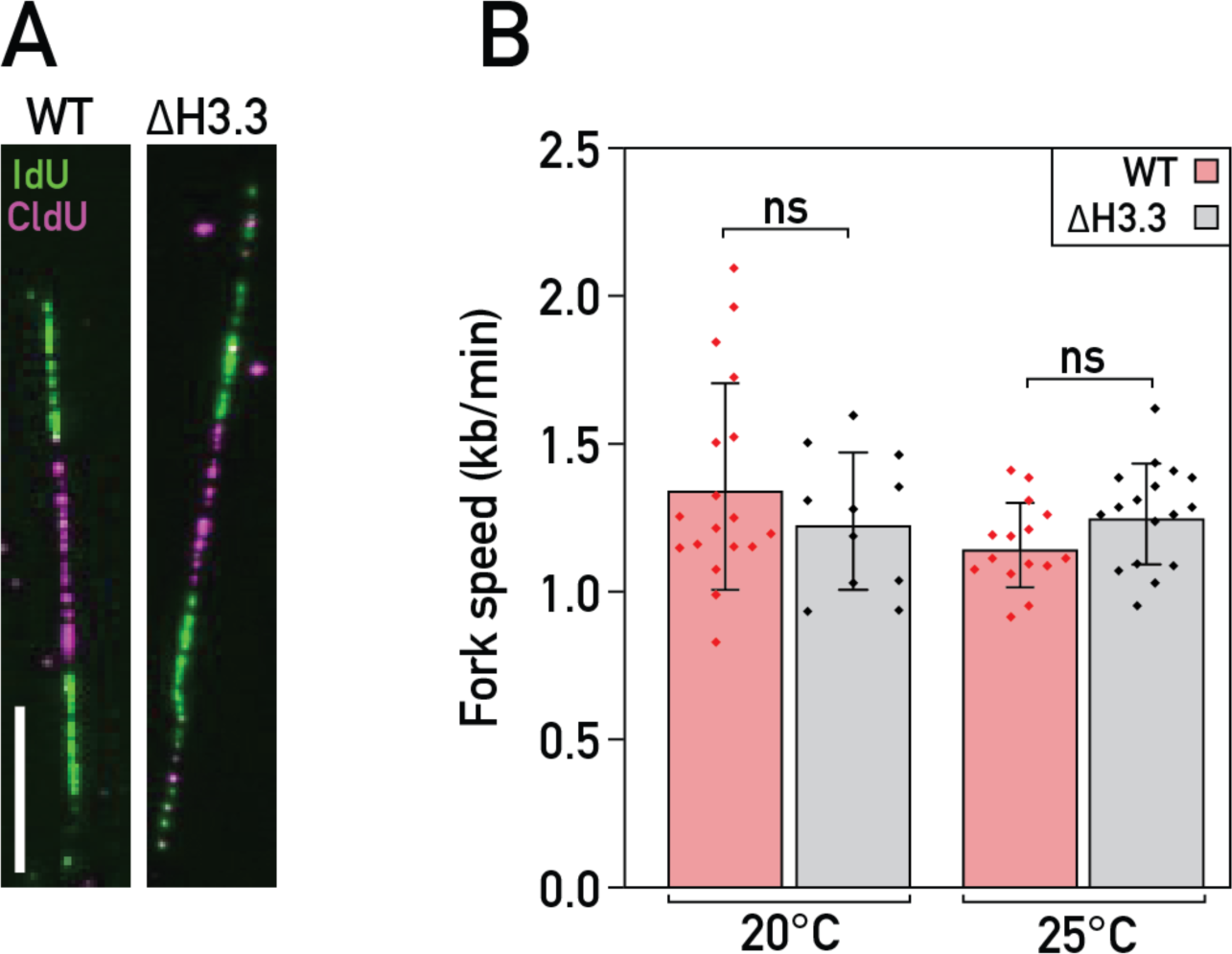
Replication fork speed is not affected by loss of H3.3.. (**A**) Representative examples of DNA combing images used to measure replication fork speed for wildtype (WT) and H3.3 null mutant (Δ H3.3). IdU incorporation is shown in green and CldU incorporation in magenta. Scale bar represents 10µm. (**B**) Fork speed determined by DNA combing at 20°C and 25°C for the WT and Δ H3.3.

### Increased DNA damage in H3.3 null mutant embryos

DNA combing assesses the speed of the moving replication fork, but it does not measure the average fork speed genome-wide, which can be affected by fork pausing, stalling or collapse. Our EdU-seq time course analysis suggested that replication origins fire normally upon loss of H3.3, but that the progression of forks is altered around origins. We therefore probed the embryos for the presence of replication stress that might be indicative of fork stalling and collapse. In *C. elegans* embryos, DNA replication defects are characterized by an increased length of the first embryonic cell cycle (Brauchle et al., 2003; Holway et al., 2006; Stevens et al., 2016). We measured cell cycle timing from the appearance of a cleavage furrow in the P0 cell to nuclear envelope breakdown of the AB cell (Figure 6A). We found that the cell cycle time was significantly increased in H3.3 null mutant embryos compared to wild type (Figure 6B). Removing the replication checkpoint through targeting the checkpoint kinase 1 (CHK-1) by RNAi restored normal cell cycle timing, confirming that the increased cell cycle length is linked to a replication defect (Figure 6B). Moreover, removal of CHK-1 resulted in an increase of the embryonic lethality in H3.3 null mutants (Figure 6C). Together, our results indicate that loss of H3.3 results in defects in DNA replication or DNA damage repair, and that these defects become detrimental upon removal of CHK-1.

**Figure 6.**
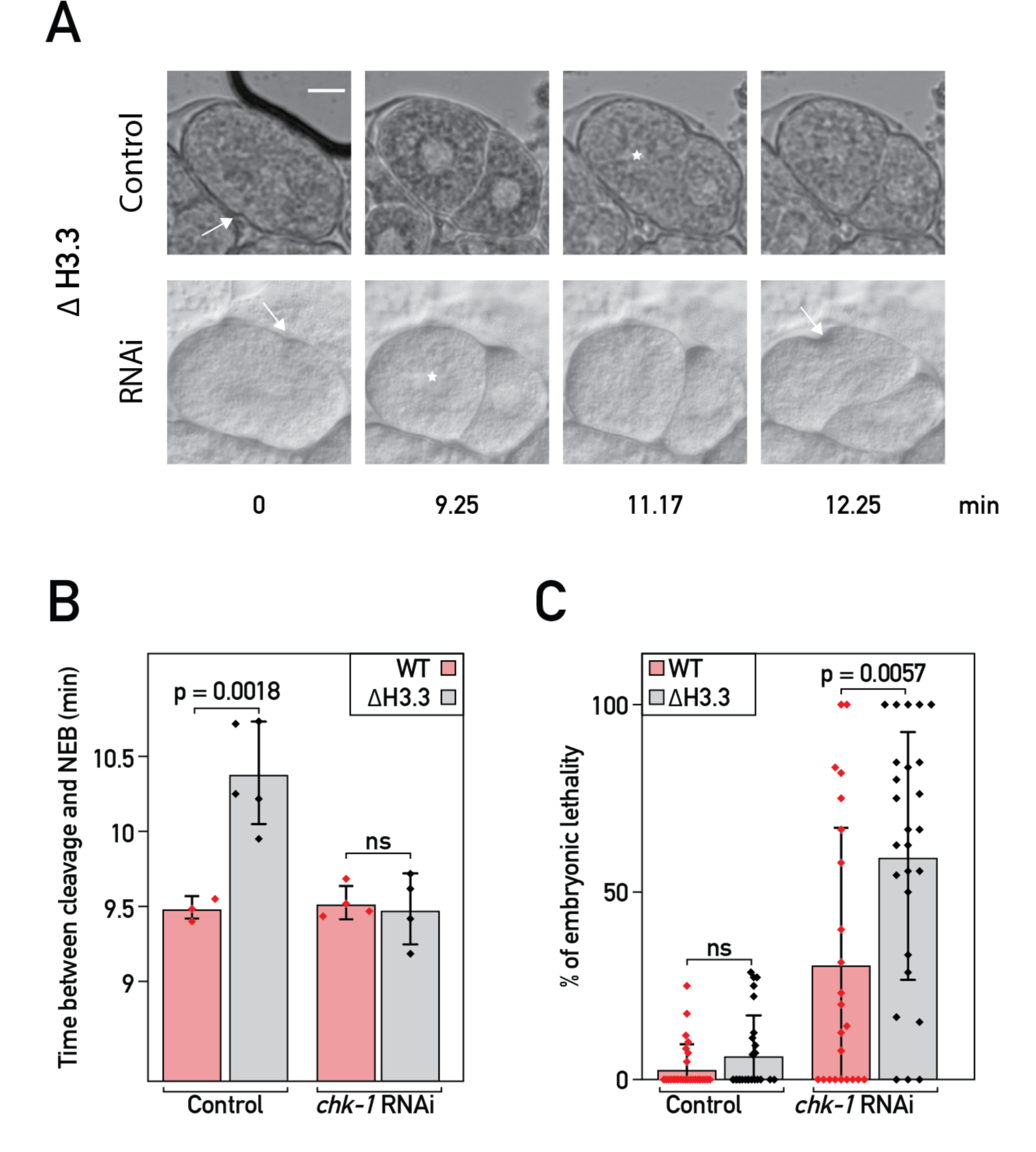
Loss of H3.3 results cell cycle delays and replication checkpoint activation. (**A**) Determination of cell cycle length, from the onset of the cleavage furrow of the first cell division (arrow) to nuclear envelope breakdown of the AB cell (star). Representative still images from a movie of a developing H3.3 null mutant (Δ H3.3) embryo with and without depletion of *chk-1* by RNAi are shown. Scale bar represents 10µm. (**B**) Cell cycle timing for wild type (WT) and Δ H3.3 worms, with and without depletion of *chk-1* by RNAi. (**C**) Embryonic lethality for WT and Δ H3.3, with and without depletion of *chk-1* by RNAi.

Taken together, we find that replication of the *C. elegans* genome is partitioned into early and late domains, and that the global timing of replication and replication origin firing is unchanged in H3.3 null mutant worms. However, loss of H3.3 leads to defective fork progression around origins and DNA replication defects at 25°C. Our results identify the replication dynamics during the embryonic cell cycle and uncover a role of H3.3 in DNA replication in *C. elegans* embryos.

## Discussion

The variant histone H3.3 has previously been linked to DNA replication, but its mechanistic role in this process, and how its function relates to its genomic localization are not well understood. In most animal models, loss of H3.3 results in lethality or sterility, making the developmental analysis of H3.3 mutant challenging (Couldrey et al., 1999; Hödl and Basler, 2009; Sakai et al., 2009; Tang et al., 2015). Our recent discovery that *C. elegans* H3.3 null mutants are viable allowed us to analyze potential developmental defects in more detail (Delaney et al., 2018). Here we describe that under stress conditions, H3.3 is required for DNA replication dynamics around replication origins and protects the genome from replication stress.

For understanding the replication defects upon loss of H3.3 in detail, we first had to develop tools to investigate DNA replication dynamics in *C. elegans* embryos. DNA replication timing in *C. elegans* had not been analyzed in detail, and replication origins have only recently been mapped by different techniques, with conflicting outcomes (Pourkarimi et al., 2016; Rodríguez-Martínez et al., 2017). We therefore adapted techniques that are based on the incorporation of EdU to determine the genome-wide replication timing (Repli-seq), and to follow the fork progression around replication origins (EdU-seq). These experiments revealed the surprising finding that despite the very short cell cycle during embryogenesis, genomic regions of early and late replication are mutually exclusive, and distributed in a non-random fashion. Partitioning of the genome into early and late replicating domains is very well described for mammalian genomes, both in cell population averages (Desprat et al., 2009; Farkash-Amar et al., 2008; Hiratani et al., 2008; MacAlpine et al., 2004) and single cells (Dileep and Gilbert, 2018; Takahashi et al., 2019) but had not yet been observed in *C. elegans*. Replication timing has been attributed to the 3D organization of chromatin into discrete domains (Marchal et al., 2019). We also observed this robust spatial organization of replication timing in *C. elegans*, despite absence of strong 3D compartmentalization (Crane et al., 2015). Chromosomes I-III are enriched for early replication domains, while most of chromosomes IV, V and X, and the telomeric regions of all chromosomes replicate late. We note that genes located in chromosomes I-III are on average more highly expressed than genes on chromosomes IV-X, and may therefore constitute an environment of open chromatin that is permissive for the binding of replication initiation factors. The domains of early replication are also enriched for H3.3. A similar correlation between early replication and H3.3 incorporation has been observed in human cells (Clément et al., 2018). Given this conserved correlation, it is maybe surprising that loss of H3.3 does not globally alter the domains of early and late replication. However, replication timing has been found to be remarkably robust, and remains unaltered in a wide range of mutants that affect chromatin organization (Marchal et al., 2019). Our results support the notion that the concentrations of some factors implicated in DNA replication are limiting, and that they will access the regions of open chromatin more readily (Mantiero et al., 2011). The concentration of replication factors is tightly controlled during S phase, and failure to export or degrade factors involved in licensing can lead to over-replication of the genome (Sonneville et al., 2012).

To comprehensively identify replication origins throughout S phase, we mapped the genomic binding sites of the origin licensing factor CDT-1 and the origin firing factor TRES-1. For determining their binding sites genome wide, we adapted the ChEC-seq method for use in *C. elegans* (Schmid et al., 2004; Zentner et al., 2015). We then combined the results with the early origins identified by EdU-seq. Our origin identification thus combines methods that detect proteins involved in origin identity with methods that rely on the identification of sites of nucleotide incorporation during origin firing. This approach allows the identification of origin firing dynamics during the cell cycle, and also includes dormant origins. *C. elegans* origins have been mapped previously using Okazaki fragment sequencing or nascent strand sequencing, both of which should identify firing origins (Pourkarimi et al., 2016; Rodríguez-Martínez et al., 2017). Our approach identified significantly fewer origins (∼1’000 vs ∼2’000 or ∼16’000, respectively), but the genomic location of the origins identified in this study largely overlap with those previously identified (Figure S2C). Previous approaches utilized the depletion of DNase *lig-1* by RNAi (Pourkarimi et al., 2016) or liquid culture (Rodríguez-Martínez et al., 2017) for the cultivation of worms for origin analysis. Both treatments result in worms that are superficially wild type, as is the case for H3.3 null mutant worms. Given our observation that even mild temperature stress can affect origin firing, and that liquid culture can affect gene expression (Çelen et al., 2018), these treatments could alter origin use and contribute to the observed differences in origin numbers. To independently estimate the number of origins required to replicate the entire genome in our system, we measured the speed of the replication fork on single molecules by DNA combing, and determined the time required to reach maximum EdU incorporation within one cell cycle. We found that a plateau of EdU incorporation is reached about 40 minutes after the first observed EdU incorporation, suggesting that cells require about 40 minutes to replicate the entire genome under these conditions (Figure S4). This is substantially longer than the S-phase duration observed in developing embryos (Sonneville et al., 2012), and we attribute the delay to the dissociation and the treatment with HU and EdU. Based on this replication time and the fork speed measured in the DNA combing experiment, we estimate the number of firing origins required for replicating the entire genome to be about 924. This number is remarkably close to the number of firing origins identified by ChEC-seq and EdU-seq (798), and we therefore think that our mapping strategy accurately reflects active replication origins. While our methods allow for the detection of origins that only fire under specific conditions, they still rely on averaging signal from very large populations of cells, and will therefore be less sensitive to detect origins that are only licensed at specific developmental time points or in specific cell types.

H3.3 is enriched at replication origins in *D. melanogaster* and *A. thaliana*, but the functional significance of this enrichment remains unclear, and no defects in origin dynamics have been detected upon loss of H3.3 (Deal et al., 2010; Eaton et al., 2011; MacAlpine et al., 2010; Paranjape and Calvi, 2016; Stroud et al., 2012). We find that H3.3 is enriched at replication origins, but consistent with the findings from *D. melanogaster*, we found no evidence that loss of H3.3 inhibits replication origin firing (Paranjape and Calvi, 2016). The EdU-seq enrichments at early origins remains unchanged, and H3.3 null mutant worms are able to activate dormant origins similar to wild type under stress conditions, as evidenced by the TRES-1 signal observed at 25°C. However, we found clear differences in EdU incorporation around origins as S phase progresses. Fork progression from origins is impaired, and the EdU-seq signal remains enriched at replication origins, in particular at those that only fire under stress conditions. The speed of the replication fork along the chromatin fiber is not affected by the loss of H3.3, as determined by DNA combing experiments. It is therefore likely that instead the frequency of fork stalling or fork collapse is increased as the fork is progressing away from the replication origins, resulting in the increased EdU-seq signal around replication origins. Our observation of the replication checkpoint activation through CHK-1 is also consistent with increased replication stress rather than a general slowing of fork progression. The defects are only observed at 25°C, and origin dynamics in wild type and H3.3 null mutant worms are indistinguishable at 20°C, consistent with the fertility defects of H3.3 null mutant worms that are only observed at higher temperatures. The reasons behind the temperature sensitivity of the replication fork upon loss of H3.3 are currently unclear.

The increased replication stress in the H3.3 null mutant could be caused directly by altering the chromatin organization, or indirectly by affecting gene expression. Decreased nucleosome density may affect replication fork progression. In *D. melanogaster* embryos, a decrease in histone concentration results in an increased cell cycle length and CHK-1 activation, similar to our observations in *C. elegans* embryos (Chari et al., 2019). However, only three genes encoding for H3.3 are expressed during embryogenesis, compared to 16 genes encoding canonical H3 (Boeck et al., 2016). It is therefore unlikely that loss of H3.3 results in a dramatic drop in nucleosome density genome wide. However, replacing H3.3 nucleosomes with H3 nucleosomes may alter the chromatin dynamics and also contribute to the observed replication stress. Absence of H3.3 may not cause replication stress directly, but may prevent fork restart after fork pausing or collapse, similar to what has been observed after DNA damage in chicken cells (Frey et al., 2014). Alternatively, loss of H3.3 could alter the transcriptional landscape genome-wide, thus resulting in increased conflicts between transcription and DNA replication. Indeed, the majority of the dormant origins, for which we find the strongest effects, are located within genes (Figure S2B). However, we previously reported that loss of H3.3 only results in minor changes in the transcriptome (Delaney et al., 2018). Loss of H3.3 could also alter the histone marks around origins. *C. elegans* replication origins were found to be enriched for H3 acetylation, H3K4 methylation and H3K27 acetylation (Pourkarimi et al., 2016), and it is possible that the presence of H3.3 aids the deposition of some of these marks. However, the causal relationship between histone modifications and the identity of replication origins is not clear. Many studies have accumulated evidence that chromatin context is important for the activity of replication origins, and we now show that H3.3 plays an important role in the fork progression following origin firing under stress condition. More research will be needed to establish a causal relationship between the chromatin context and replication origin formation, to understand how the timing of origin firing is regulated, and to determine how the replication fork interacts with nucleosomes containing different histone modifications or variants.

## Materials and Methods

### Worm culture and strain generation

Worms were grown on NGM plates seeded with OP50 for maintenance, and on peptone-rich NGM plates seeded with NA22 for large-scale experiments. The sequence coding for MNase was inserted at the *cdt-1* and *tres-1* loci using CRISPR/Cas9 as described in (Arribere et al., 2014), using plasmids with 1 kb homology arms as repair templates. A list of the strains used in this study is given in Table S1. RNAi experiments were carried out by feeding, using clones from the Ahringer library (Kamath et al., 2003). For experiments measuring effects at 25°C, worms were grown at this temperature for at least 5 generations.

### Phenotype analysis

For determining embryonic lethality at 25°C, L4 worms were placed on plates containing RNAi or control food overnight. Single adults were transferred to new plates for 3h and then removed. The number of eggs, hatched larvae and adults was counted on subsequent days to determine the penetrance of embryonic lethality. Plates with 0 or 1 embryos were excluded from the final count. Data from three independent experiments with 10 plates per experiment and condition were used. Total number of embryos: WT control = 445, Δ H3.3 control = 240, WT *chk-1* RNAi = 270, Δ H3.3 *chk-1* RNAi = 251.

For determining the cell cycle length during early embryonic cell divisions at 25°C, L4 worms were placed on plates containing RNAi or control food overnight. Adults were cut in egg buffer (118mM NaCl, 48 mM KCL, 2mM CaCl2, 2mM MgCl2 and 25mM Hepes pH7,5) to release young embryos. These embryos were mounted on 2% agarose pads and recorded every 10 s with DIC settings on a Leica DMI8 microscope, with the temperature maintained at 25°C. Cell cycle length was measured from the appearance of the cleavage furrow of the P0 cell to nuclear envelope breakdown of the AB cell. WT control = 3, Δ H3.3 control = 5, WT RNAi = 4, Δ H3.3 RNAi = 4.

For determining the subcellular localization of CDT-1 and TRES-1, gravid adult worms were washed in PBS containing 0.1% Triton X-100, anesthetized in DB buffer (50 mM sucrose, 75 mM HEPES pH 6.5, 60 mM NaCl, 5 mM KCl, 2 mM MgCl_2_, 10 mM EGTA pH 7.5, 0.1% NaN_3_) and cut in half to release embryos. The embryos were transferred to poly-L-lysine-coated slides, freeze-cracked and fixed in cold methanol for 5 min. Slides were washed two times in PBS and incubated with anti-HA antibody (mAb 42F13, 1:60) for CDT-1 staining or FLAG antibody (Sigma-Aldrich, F7425-.2MG, 1:200) for TRES-1 staining, overnight at 4°C. After two washes in PBS, slides were incubated with secondary antibodies (Alexa Fluor 488 goat anti-mouse or Alexa Fluor 488 goat anti-rabbit; 1:700) for 1 h at room temperature, washed once in PBS and counterstained with DAPI (2 µg/ml) for 15 minutes. After a final PBS wash, the slides were mounted in VECTASHIELD Antifade Mounting Medium. Images were acquired with Leica SP8 confocal microscope. For all the images Z-sections with 0.3 µm step were acquired, and processed with Fiji software (maximum Z-projection, contrast adjustment, Gaussian blur filter r=0.5) (Schindelin et al., 2012).

### Chromatin Endogenous Cleavage (ChEC)

Worm cultures were synchronized by embryo isolation with sodium hypochlorite for 4 generations. Embryos were treated with chitinase (1h, RT, 2 U/ml), washed twice in buffer A (15 mM Tris pH 7.5, 0.1 mM EGTA, 0.34 M sucrose, 0.2 mM spermine, 0.5 mM spermidine, 0.5 mM PMSF) and then resuspend in buffer A supplemented with detergents (0.25% NP-40 and 0.1% Tx-100). Nuclei were released with a glass dounce homogenizer and 15 stokes using the loose inserting pestle and 15 stokes using the tight inserting pestle. Debris were spun down at 100 g for 2 minutes and the supernatant, containing nuclei, was saved. The debris were resuspended in buffer A supplemented with detergents and the nuclei isolation step was repeated. The supernatants containing the nuclei were pooled, and the nuclei were pelleted at 1000 g for 10 min. Nuclei were washed once in buffer A, once in buffer B (15 mM Tris pH 7.5, 80 mM KCl, 0.1 mM EGTA, 0.2 mM spermine, 0.5 mM spermidine, 1 mM PMSF), and resuspended in buffer B. MNase was activated by the addition of CaCl_2_ to a final concentration of 2 mM at 30°C. Samples were incubated for 5 to 10 minutes (for CDT-1 ChEC), 5 to 7 minutes (for TRES-1 ChEC) or for 1 h (free MNase control). For the control sample, 0.3 U/ml of MNase (Biolabs, Cat. No. M0247S) was added at the same time as the CaCl_2_. MNase was inactivated by adding an equal volume of 2x stop buffer (400 mM NaCl, 20 mM EGTA). Samples were treated overnight with Proteinase K and SDS at 55°C. Phenol-chloroform-isoamyl treatment was used to remove Proteinase K prior the RNase treatment (Roche, Cat. No. 11119915001) for 1h30 minutes at 37°C. After chloroform extraction, DNA was precipitated overnight with ethanol and 0.25 M NaCl. DNA was resuspended overnight in TE prior to constructing the libraries using NEBNext® Ultra™ II DNA Library Prep. Fifty base pair paired-end read sequencing reactions were then performed on an Illumina Hi-Seq 2500 sequencer at the Genomics Platform of the University of Geneva, except for TRES-1 ChEC-seq samples from 25°C, which were sequenced with one hundred base pair single-end read sequencing reactions.

### Embryonic cell extraction for EdU incorporation or DNA combing

Cells were extracted from embryos as described in (Bianchi and Driscoll, 2006). Briefly, embryos were isolated using sodium hypochlorite, treated with chitinase (1h, RT, 2U/ml), washed once in M9 and resuspend in 10ml of L-15 medium (supplemented with 10% FBS (heat inactivated, Gibco), 41 mM sucrose and 1% pen/strep (1:100)). Cells were dissociated with three passes through a syringe with a 22 gauge needle and two passes through a syringe with a 26 gauge needle. Debris were pelleted by spinning at 80 g for 1 min, and the supernatant, containing cells, was saved. Debris were resuspended in supplemented L-15 medium and cell extraction was repeated twice more. Supernatants were pooled together and filtered using 5 micron filters. Cells were pelleted at 500 g for 15 min and resuspended in supplemented L-15 medium.

### EdU-seq

EdU-seq was carried out as described (Macheret and Halazonetis, 2019) with minor modifications. Embryonic cells were synchronized by addition of 20 mM of HU (Sigma, Cat. No. H8627) during 1h at 20°C or 25°C. Cells were washed twice with PBS to remove HU and resuspended in L-15 medium. At desired time points, 25 µM of EdU and 20 mM of HU were added for 10 minutes. Cells were washed twice with PBS, fixed with 90% methanol and stored at −20°C. Cells were washed with ice-cold PBS and permeabilized with PBS containing 0.2% Triton X-100 for 30 minutes at room temperature in the dark. EdU was coupled to a cleavable biotin-azide linker (Azide-PEG(3+3)-S-S-biotin) (Jena Biosciences, Cat. No. CLK-A2112-10) using the reagents of the Click-it Kit (Invitrogen, Cat. No. C-10424). Cells were washed with PBS and treated overnight at 50°C with proteinase K in the lysis buffer. DNA was extracted by phenol/chloroform/isoamyl extraction followed by chloroform extraction. DNA was precipitated overnight at −20°C with presence of ethanol and 0.25 M NaCl and resuspended overnight in TE. For an estimation of the time required to replicate the entire genome, cells were isolated, synchronized using HU, and released in the presence of EdU as described above. Cell aliquots were harvested between 0 and 130 min. Cell permeabilization and click-it reaction were done as described above to couple Alexa Fluor™ 647 Azide (Thermofisher Cat. No. A10277) to the EdU-labeled DNA. Cells were washed twice with PBS and treated with RNase (Roche, Cat. No. 11119915001) for 30 minutes at 37°C. PI (Sigma, Cat. No. 81845) was added, and samples incubated overnight at room temperature in the dark. Levels of EdU incorporation were assessed by flow cytometry (Gallios, Beckman Coulter).

### Repli-seq

Asynchronous embryonic cells extracted from worms grown at 25°C were incubated for 5 minutes with 25 µM EdU (Invitrogen, Cat. No. A10044), washed twice with PBS and fixed with 90% methanol overnight. After fixation and click-it reaction as described in the EdU-seq section, cells were washed twice with PBS and treated with RNase (Roche, Cat. No. 11119915001) for 30 minutes at 37°C. Propidium iodide (Sigma, Cat. No. 81845) was added, and samples were incubated overnight at room temperature in the dark. Cells were sorted according to their DNA content using a MoFlo Astrios flow sorter (Beckman Coulter) at the Flow Cytometry platform of the Medical Faculty of the University of Geneva. DNA was extracted as described in the EdU-seq section.

### Isolation and sequencing of EdU-labelled DNA from EdU-seq and Repli-seq

Five to eight µg of DNA were sonicated with a bioruptor sonicator (Diagenode) to obtain fragments of 100-500 bp. EdU-labelled fragments were isolated using Dynabeads MyOne Streptavidine C1 (Invitrogen, Cat. No. 65001) as previously described (Macheret and Halazonetis, 2019). DNA was eluted by the addition of 2% of B-mercaptoethanol (Sigma, Cat. No. M6250) and incubation for 1h at room temperature. The eluted DNA, as well as fragmented total DNA, was used for library preparation using the TruSeq ChIP Sample Prep Kit (Illumina, Cat. No. IP-202-1012). One hundred base pair single-end read sequencing reactions were performed on an Illumina Hi-Seq 2500 sequencer at by the Genomics Platform of the University of Geneva.

### Data processing, domain annotation and peak calling

Sequencing reads were mapped to the *C. elegans* reference genome WBcel215 using Novoalign software (default parameters, producing SAM format files). Reads were transformed into 1’000 bins size (ChEC-seq and EdU-seq) or 10’000 bins size (Repli-seq) (JSON format) with a custom script. Biological replicates that passed the quality control were merged (Repli-seq: WT early (2), WT mid (1), WT late (3), Δ H3.3 early (2), Δ H3.3 mid (2), Δ H3.3 late (2), ChEC-seq: CDT-1 20°C (5), TRES-1 20°C (4), TRES-1 25°C (3), TRES-1 25°C Δ H3.3 (3), EdU-seq: WT 0 min 20°C (4), all the other EdU-seq data (2)). A custom R script was used to transform JSON files into bedgraphs. All files were normalized to the same number of reads. Experimental samples were then normalized by subtracting the number of reads present in the corresponding bin in the control file. The control samples were DNA from nuclei treated with Free MNase (ChEC-seq), genomic DNA from cells treated with HU (EdU-seq), or genomic DNA from cells without treatment (Repli-seq). Finally, the data were smoothed with a sliding window averaging 5 consecutive bins using a custom R script. Genome Browser images were obtained by using the Sushi Bioconductor package in R.

Domains of early and late replication were defined according to the signal enrichment present in early and late Repli-seq in WT samples. Bins with positive signal in early and negative signal in late datasets were defined as early domains, and vice versa for the ones with negative signal in early and positive signal in late as late domain. The bins with either positive signal in early and late repli-seq or negative signal in both datasets were not assigned to a domain. Domains smaller than 5kb were merged with the previous domain.

For peak calling, normalized and smoothed data was converted to SGA files using the “ChIP-convert” tool in the Vital-it platform (https://ccg.epfl.ch//chipseq/). The output SGA files were used as input for peak calling with the tool “ChIP-peak” of the Vital-it platform with these specific parameters: Cut-off = 1000, Read counts = 10, Window = 3000, Vicinity = 5000, strand = any, Repeat Masker = on, Refine peak position = off, Relative Enrichment 325 (CDT-1), 450 (TRES-1) or 350 (EdU). Peak lists from the ChEC-seq and EdU-seq experiments were merged, and duplicate peaks removed. Peaks are considered duplicate if two datasets contains a peak within the same 5kb window. To avoid sequencing artefacts, peaks that were also enriched in the genomic DNA sample were removed. The final list of peaks was then clustered in 15 using deepTools (Ramírez et al., 2016). Clusters with similar enrichment were merged together and one cluster, containing signal only in the EdU-seq sample, was deleted, as the sites in this cluster did not show fork movement after release from the HU block. Heatmaps were created using deepTools (Ramírez et al., 2016). The genomic annotation of early, late and dormant origins was determined using PAVIS website (https://manticore.niehs.nih.gov/pavis2/) (Huang et al., 2013). Sequencing data has been deposited at the Gene Expression Omnibus (GEO) under accession number GSE140804.

### DNA combing and staining

DNA combing was performed as described in (Michalet et al., 1997). Embryonic cells were incubated with 20 mM HU for 1h at 20°C or 25°C, washed twice with ice-cold PBS, and incubated with 40 µM CldU for 20 minutes at the desired temperature. Cells were washed twice with L-15 and then incubated in presence of 400 µM IdU for 20 minutes at the desired temperature. Cells were washed with PBS and frozen at −20°C. Cells were embedded into agarose plugs, and DNA was processed using the FiberPrep® DNA extraction kit (Genomic Vision). The combing and antibody incubation was done as described in (Costantino et al., 2014). The DNA combing slides were imaged on a Leica DM5000B microscope, and the images were analyzed with Fiji software (Schindelin et al., 2012). The analysis was carried out with data from six independent experiments with a total of 49 measurements (WT, 20°C), 32 measurements (WT, 25°C), 28 measurements (Δ H3.3, 20°C) and 49 measurements (Δ H3.3, 25°C).

## Acknowledgements

We thank Morgane Macheret and Thanos Halazonetis for sharing unpublished protocols, Jonathan Mailler and Laura Padayachy for discussion and help with the DNA combing experiments, Dario Menendez for help with strain generation, Caroline Gabus for reagent preparation, Nicolas Roggli for help with data analysis and image processing, Florian Huber and Ioannis Xenarios for help with the data analysis and Uli Laemmli and Gabriel Zentner for reagents and advice when establishing the ChEC method. We thank all the Steiner lab members for discussion and comments on the manuscript. We thank Thanos Halazonetis, Damien Hermand, Jonathan Mailler, Laura Padayachy, Reinier Prosee, David Shore and Maksym Shyian for comments on the manuscript. Some strains were provided by the CGC, which is funded by NIH Office of Research Infrastructure Programs (P40 OD010440). We are grateful to the Genomics Platform of the iGE3, the Bioimaging Center of the Faculty of Sciences and the Flow Cytometry platform of the Medical Faculty at the University of Geneva. The work was funded by the Swiss National Science Foundation (Grants 31003A_156774 and 31003A_175606) and the Republic and Canton of Geneva.

## Author contributions

FAS and MS conceived the study, MS carried out genetics, genomics and DNA combing experiments, JMW carried out staining experiments, MS analyzed the data, MS and FAS prepared the manuscript, FAS acquired funding.

---

**Figure S1.**
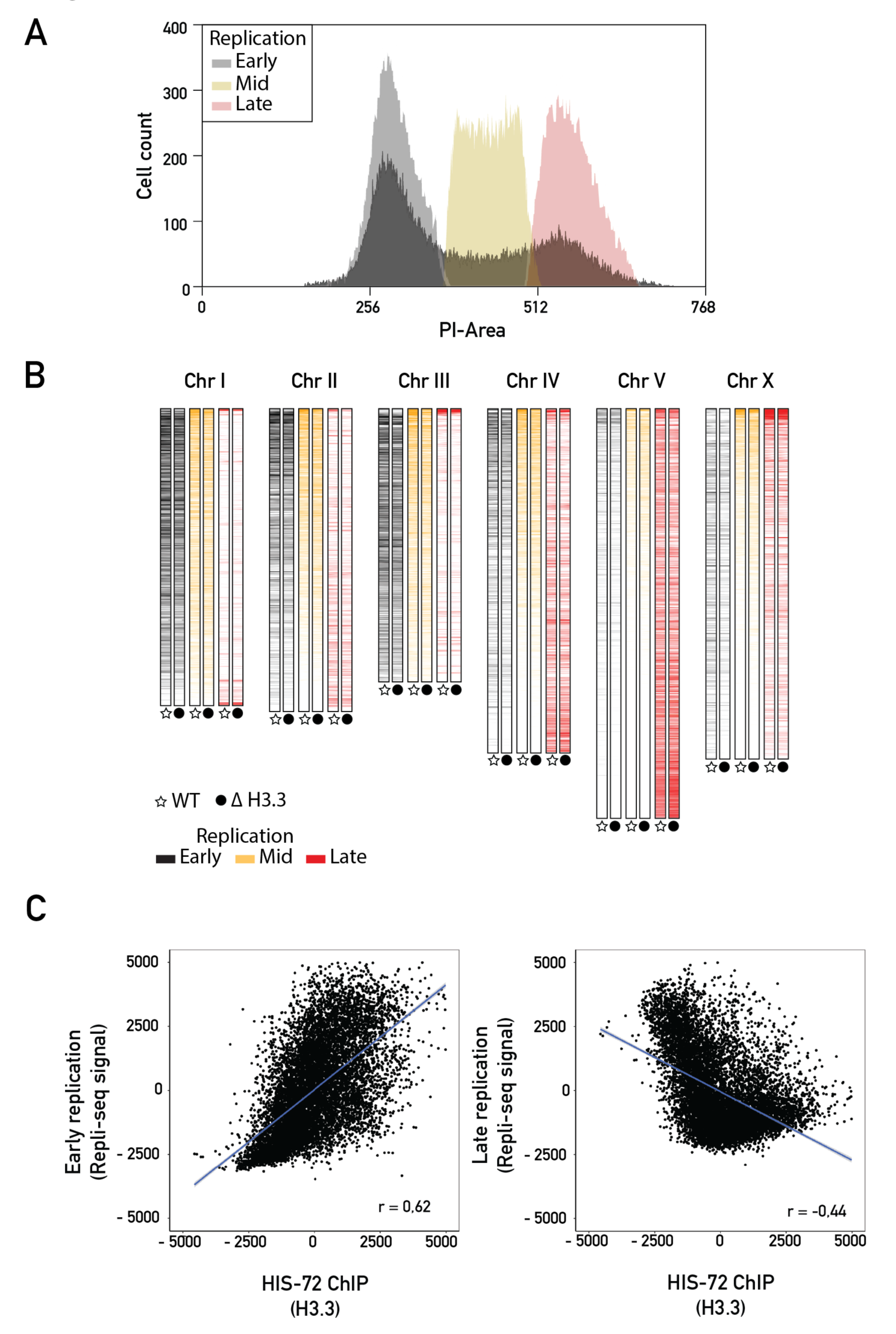
Characteristics of early and late replicating domains. (**A**) Boundaries used to sort embryonic cells according to their DNA content. Plot showing the number of cells counted (cell counts) as a function of the amount of PI incorporation (PI-Area). Cells were sorted according to their DNA content into early replication (gray), mid replication (yellow) and late replication (red). (**B**) Color-coded replication timing for each chromosome. Repli-seq signal from early (black), mid (orange) and late (red) S phase for wild type (WT) and H3.3 null mutant (Δ H3.3) worms. Data is shown according to the chromosomal location. (**C**) Correlation plots between Repli-seq data for early (left panel) or late (right panel) replication with HIS-72 ChIP-seq data (Delaney et al., 2019), in 10kb bins. The Spearman’s rank correlation coefficient (r) is shown.

**Figure S2.**
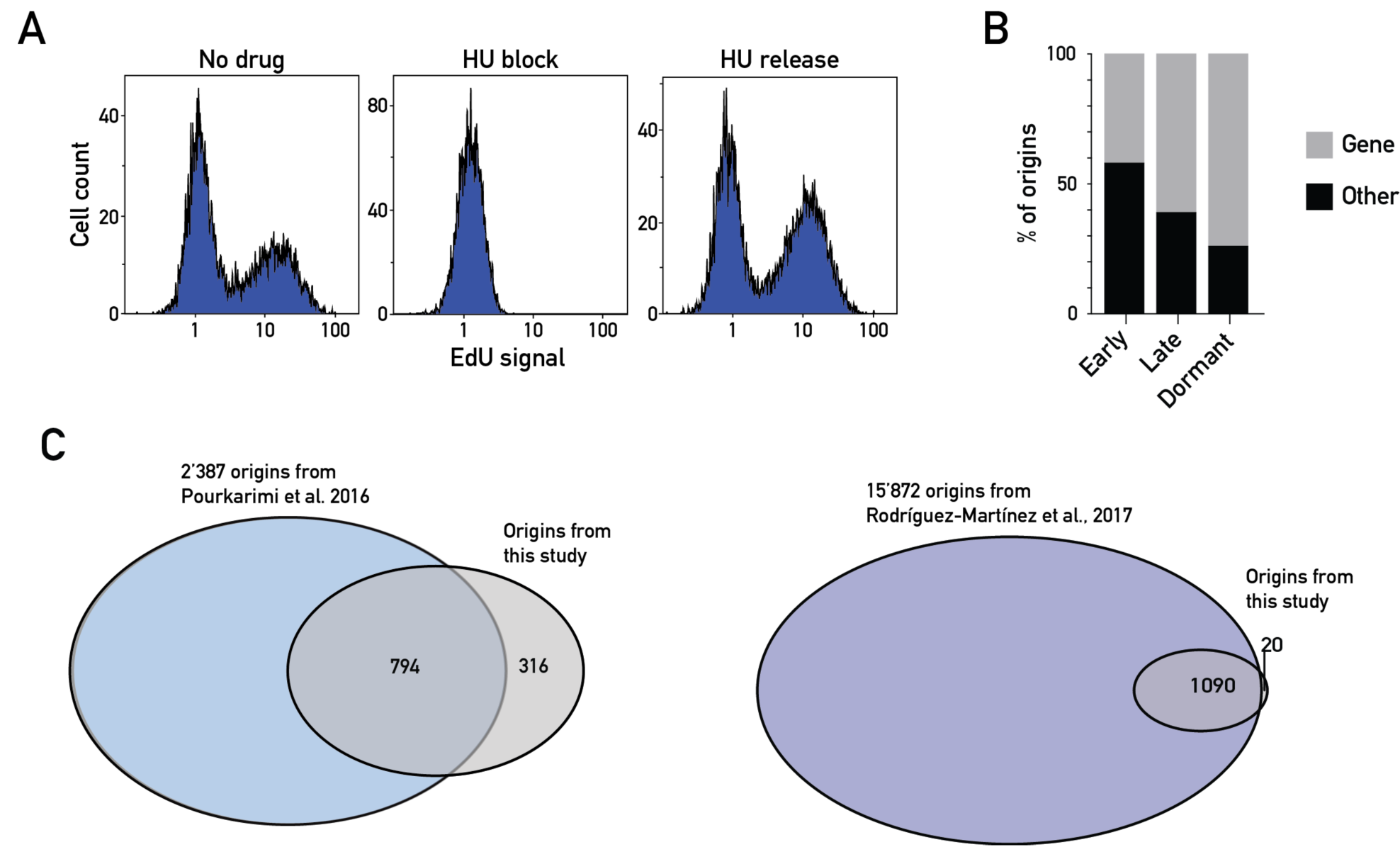
Characteristics of replication origins. (**A**) FACS profiles from HU block experiments for EdU-seq. Plots show distribution of EdU incorporation across cell populations in absence of HU (left panel), upon HU block (center panel) and 30 minutes after release from the HU block (right panel). (**B**) Classification of early, late and dormant origins found in this study based on genomic location within (genic) or outside (non-genic) gene-coding regions. (**C**) Venn diagrams representing the number of origins identified in this study that overlap with the origins found in (Pourkarimi et al., 2016) (left) and in (Rodríguez-Martínez et al., 2017) (right).

**Figure S3.**
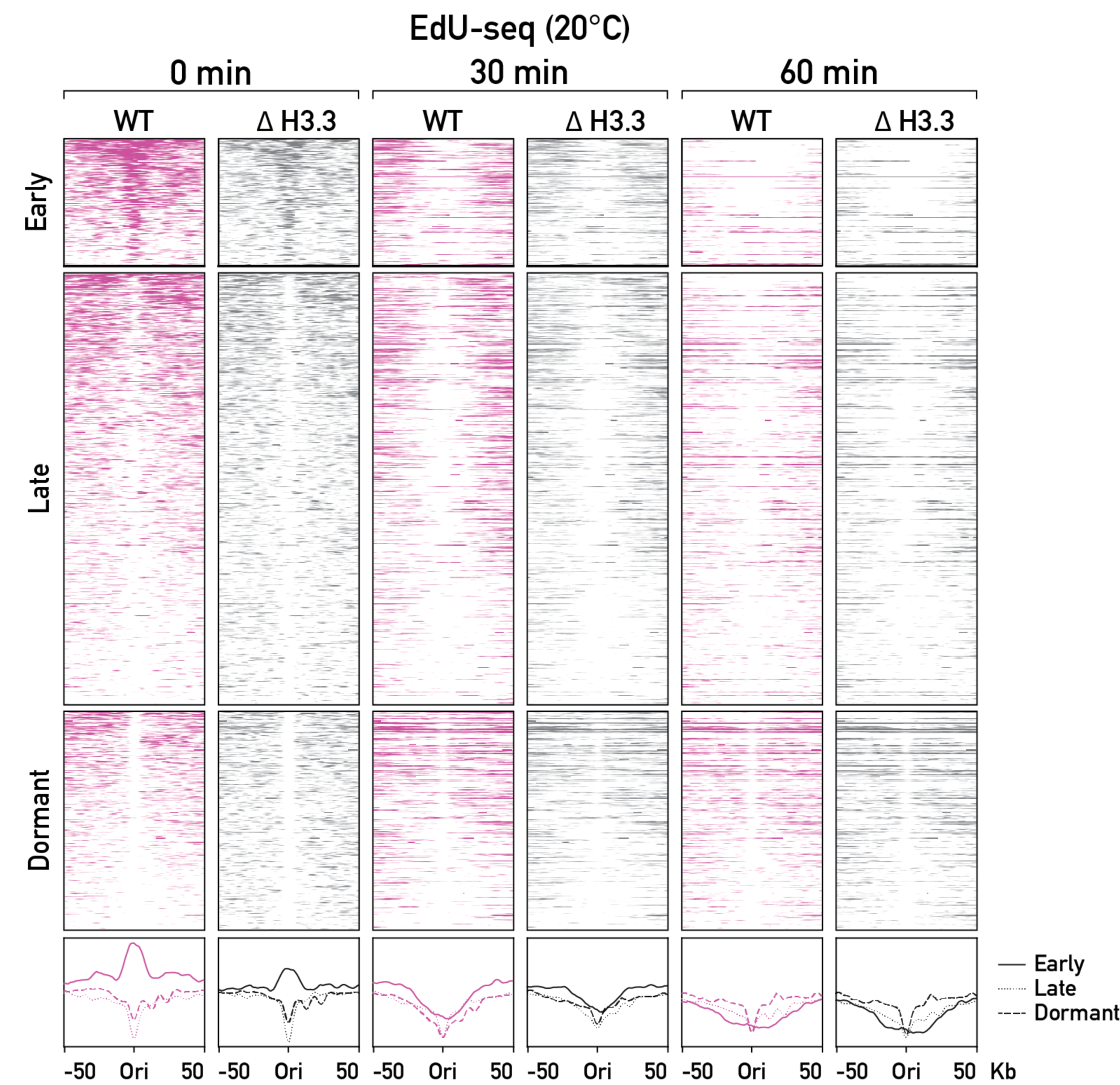
Replication origin dynamics under non-stress conditions. EdU-seq time course, for wild type (WT, pink) and H3.3 null mutant (Δ H3.3, gray) worms grown at 20°C. Heatmaps (top) and average plots (bottom) of EdU-seq signal at 0, 30 and 60 minutes, at all origins. Signal is shown for a 50 kb window around each origin.

**Figure S4.**
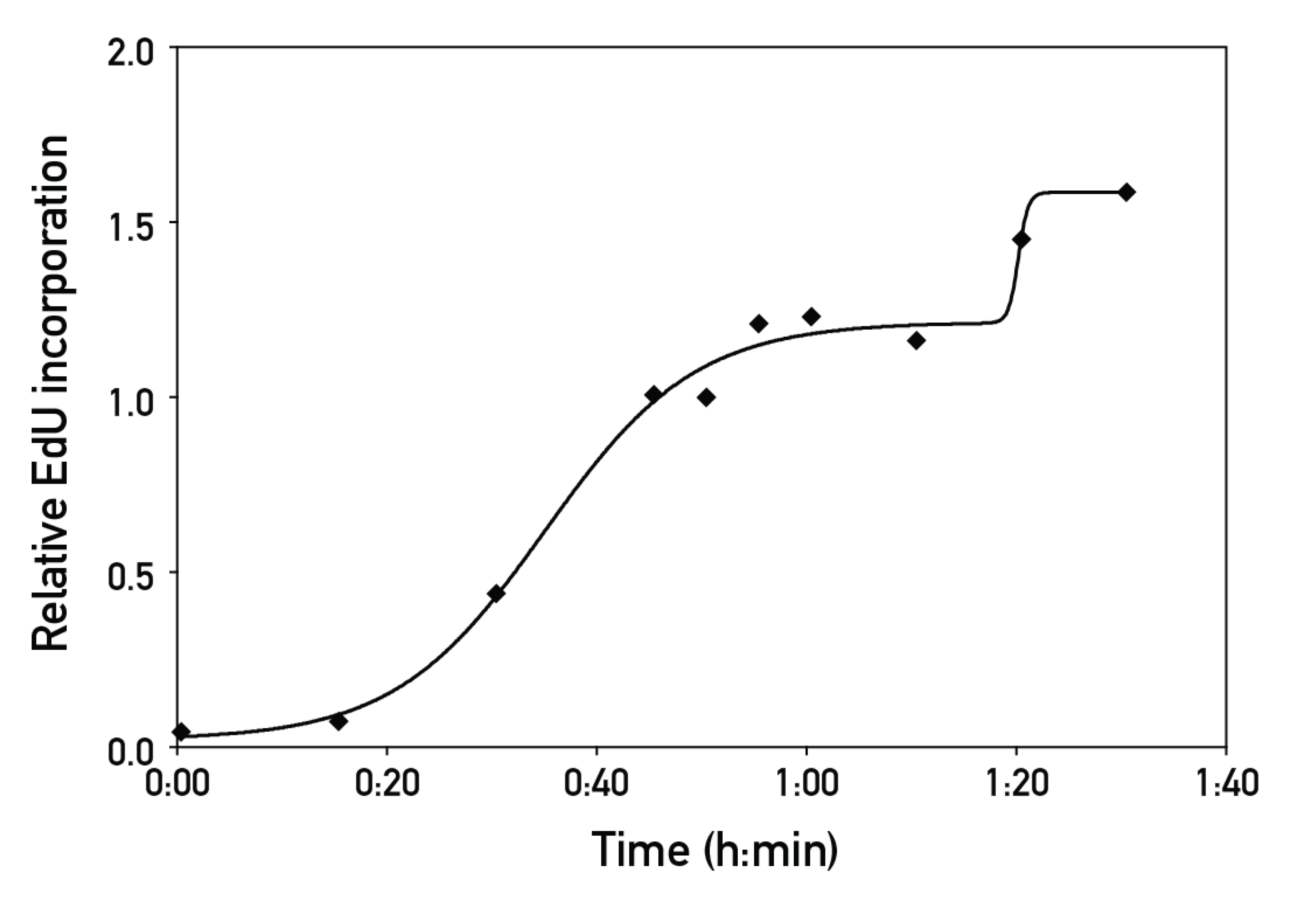
Estimation of S-phase duration. Time course of EdU incorporation after HU release at 20°C. Relative EdU incorporation is shown as a function of the time of incubation in presence of EdU. The curve was obtained using a bell-shaped model with GraphPad Prism version 8.0.0.

**Table S1.** Overview of *C. elegans* strains used in this study.

| Strain | Genotype | Source | Description |
| --- | --- | --- | --- |
| N2 |  | CGC | Wild type |
| FAS43 | *his-69&his-70(uge44) III; his-72(tm2066) III; his-74(uge18) V; his-71(ok2289) X* | Delaney et al., 2018 | H3.3 null mutant |
| FAS50 | *cdt-1(uge34 [MNase::cdt-1]) I* | This study | CDT-1 ChEC in WT background |
| FAS137 | *tres-1(uge102[MNase::tres-1]) I* | This study | TRES-1 ChEC in WT background |
| FAS154 | *tres-1(uge104[MNase::tres-1]) I; his-69&his-70(uge44) III; his-72(tm2066) III; his-74(uge18) V; his-71(ok2289) X* | This study | TRES-1 ChEC in ΔH3.3 background |

